# Topographically localised modulation of tectal cell spatial tuning by natural scenes

**DOI:** 10.1101/2021.12.03.471079

**Authors:** Thomas Trevelyan James Sainsbury, Giovanni Diana, Martin Patrick Meyer

## Abstract

Visual neurons can have their tuning properties contextually modulated by the presence of visual stimuli in the area surrounding their receptive field, especially when that stimuli contains natural features. However, stimuli presented in specific egocentric locations may have greater behavioural relevance, raising the possibility that the extent of contextual modulation may vary with position in visual space. To explore this possibility we utilised the small size and optical transparency of the larval zebrafish to describe the form and spatial arrangement of contextually modulated cells throughout an entire tectal hemisphere. We found that the spatial tuning of tectal neurons to a prey-like stimulus sharpens when the stimulus is presented in the context of a naturalistic visual scene. These neurons are confined to a spatially restricted region of the tectum and have receptive fields centred within a region of visual space in which the presence of prey preferentially triggers hunting behaviour. Our results demonstrate that circuits that support behaviourally relevant modulation of tectal neurons are not uniformly distributed. These findings add to the growing body of evidence that the tectum shows regional adaptations for behaviour.

## Introduction

Natural visual scenes are complex, requiring animals in the wild to localise and identify salient visual features such as potential predators or prey that may be masked by other objects or textured backgrounds. One neural mechanism that is thought to be important for processing natural scenes is contextual modulation (***Krause and Pack*** (***2014***)). Here the firing properties of a neuron responding to a stimulus within its receptive field (RF) can be modulated by stimuli outside it (nRF) (***Allman et al***. (***1985***); ***Levitt and Lund*** (***1997***); ***Angelucci et al***. (***2002***)). Contextual modulation has be shown to affect tuning to size (***Barlow*** (***1953***); ***Hartline et al***. (***1956***)), contrast (***Levitt and Lund*** (***1997***)), orientation (***Okamoto et al***. (***2009***)) and for discriminating local motion (***Sun et al***. (***2002***, ***2006***)). Furthermore, contextual modulation has been implicated in figure-ground separation (***Allman et al***. (***1985***)), detexturisation (***Gheorghiu et al***. (***2014***)), generating “pop-out” phenomena (***Schmid and Victor*** (***2014***); ***Zhaoping and Zhe*** (***2012***); ***Zhaoping*** (***2008***); ***Ben-Tov et al***. (***2015***); ***Knierim and Essen*** (***1992***)) and sparsifying population activity that enhance coding efficiency (***Vinje and Gallant*** (***2000***, ***2002***); ***Haider et al***. (***2010***)). Importantly, recent studies in have highlighted that these effects are most prominent when the nRF contains naturalistic features and that the circuits that implement contextual modulation in mice require visual experience to develop (***Guo et al***. (***2005***); ***Pecka et al***. (***2014***)). Therefore, it has been suggested that contextual modulation in the visual system is integral for processing natural scenes and that it is itself shaped by the statistics of natural scenes during development.

However, presenting stimuli at different positions within egocentric visual space can trigger different visually driven behaviours. This often reflects non-uniform mapping of specific cell types throughout the visual system, generating regional specialisation within the visual field (***Zimmermann et al***. (***2018***); ***Zhang et al***. (***2012***); ***Avitan et al***. (***2019***); ***Förster et al***. (***2020***)). For example, larval zebrafish are most likely to orientate itself towards prey when the prey is located 40 degrees lateral to the fishes heading direction when compeared to the rest of the visual field (***Romano et al***. (***2015***); ***McElligott and O’Malley*** (***2005***); ***Lagogiannis et al***. (***2019***)). This raises the question of whether the degree of contextual modulation also varies across the topographic axes of visual areas of the brain. In this study, we take advantage of the optical transparency and small size of the larval zebrafish brain to examine how naturalistic visual scenes modulate the responses of tectal neurons to prey-like stimuli and how, if present, such modulation changes throughout the tectal volume. Strikingly, we find that such modulation occurs within a spatially restricted region tectum. This region represents a point of visual space in which the presence of prey preferentially triggers hunting routines. Furthermore, we show that the circuits that support such contextual modulation do not require sensory experience for their proper development. Our results demonstrate that tectal circuits that support contextual modulation are localised in behaviourally relevant topographical regions.

## Results

To determine if responses of neurons in the zebrafish tectum are modulated according to the context of the visual scene, the optic tectum of 7 day post-fertilisation (dpf) larvae were imaged using 2-photon volumetric imaging whilst stimuli were presented to the contralateral (right) eye. The stimuli consisted of prey-like stimuli (moving 5° black dots), that were displayed in a pseudorandom order at 7 different locations along visual azimuth. These stimuli were displayed in two blocks that differed in their backgrounds. In one block the background contained naturalistic features in the form of a picture of gravel (textured block) whereas the other was a grey screen (grey block). Each moving spot was presented 7 times at each of the locations in each block. We then examined how tuning to stimulus location (spatial tuning) was modulated by context (background) (Fig. 1A-E).

**Figure 1.**
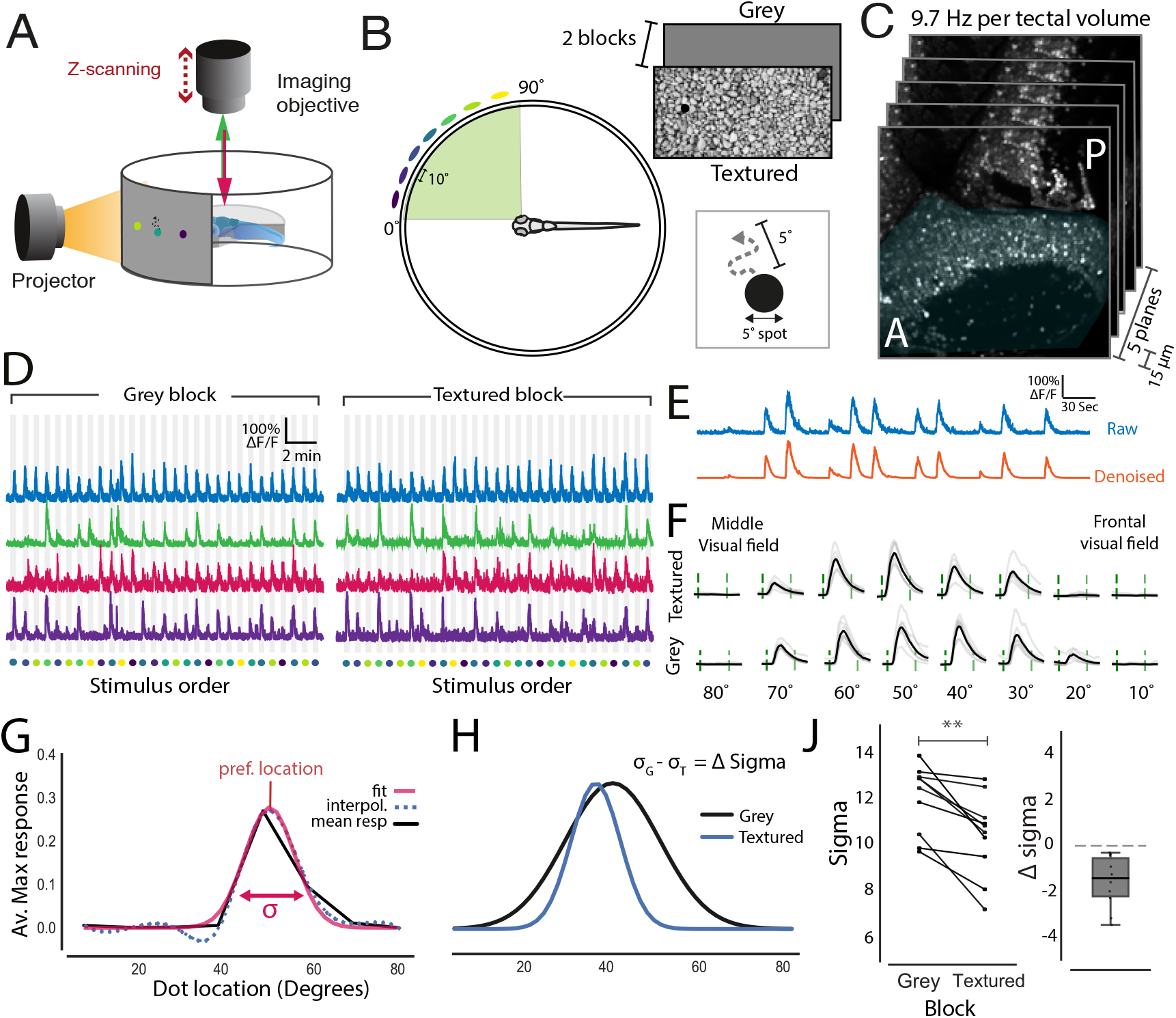
Presenting stimuli over a textured background sharpens the spatial tuning of tectal neurons. (A)Schematic of the imaging setup where larvae were head fixed in agarose allowing for visual stimuli to be projected onto a semi-cylindrical screen whilst neural activity is monitored via 2-photon volumetric imaging. (B)5° moving dots (virtual prey) were presented at 7 different locations along visual azimuth separated by 10°. These dots moved randomly within a 5° degree neighboured and were presented in two blocks with different backgrounds-grey screen or textured (gravel). (C) Imaging volumes of the contralateral tectum (shaded blue) consisted of 5 optical sections that were taken 15*μ*m apart at an imaging speed of 9.7 Hz per volume. (D) Normalised fluorescence traces from cells that are active in both the grey and textured blocks. Stimulus location is colour-coded as in B. (E) Raw fluorescence traces were denoised to generate smoothed calcium signals. (F) Mean responses to each of the stimulus locations (black) and individual repetitions (grey) for an example cell. Stimulus epoch start and end are indicated by the green dashed lines. (G) Spatial tuning curves for each cell were calculated by fitting a Gaussian to the interpolated average maximum response to each stimulus location. The preferred stimulus location was taken as the peak and tuning sharpness was taken as the standard deviation - sigma) of the Gaussian fit. (H) Tuning fits for an example neuron for both the textured and grey backgrounds. From this a neurons change in sigma (Δ sigma) can be calulated by subtracting its sigma value for the grey background (*σ_G_*) from its sigma for the textured background (*σ_T_*). (I) Left: mean sigma values for tectal neurons in both textured and grey blocks. Each connected line represents one fish (n = 10). Sigma was reduced for all fish in the textured block relative to the grey block (Paired T-test, p=.002). Right: A box plot showing the mean change in sigma for each fish (Δ sigma) between the two blocks. Dotted line indicates zero change. ** = p<.01.

Spatial tuning curves for each neuron were calculated by fitting the average responses to each stimulus location with a Gaussian (Fig. 1F-G). Tuning width, defined as the standard deviation of the Gaussian fit, could then be compared between blocks. This revealed that for all imaged fish (n=10) the average tuning width was reduced when stimuli were viewed in the textured block relative to the grey block (Fig. 1H). Furthermore, the mean change in sigma (Δ sigma) for all fish was negative, with a mean reduction in sigma of −1.5. This suggests that the spatial tuning of tectal neurons to prey-like stimuli is sharpened when viewed within complex and naturalistic visual scenes.

To examine whether contextual modulation occurs uniformly along visual azimuth, sigma values for neurons in both blocks were plotted against their preferred stimulus location. This revealed that neurons with preferred tuning location of 35-50° demonstrated significantly reduced sigma values (Fig. 2A). Importantly, this area in visual space is where small orientating movements towards prey, known as “J-turns”, are most likely to be triggered (Fig. 2C-D) (**?**). This suggests that these neurons may be important for identifying the position of local stimuli, such as prey, within complex natural scenes and that this information may be important for driving j-turns.

**Figure 2.**
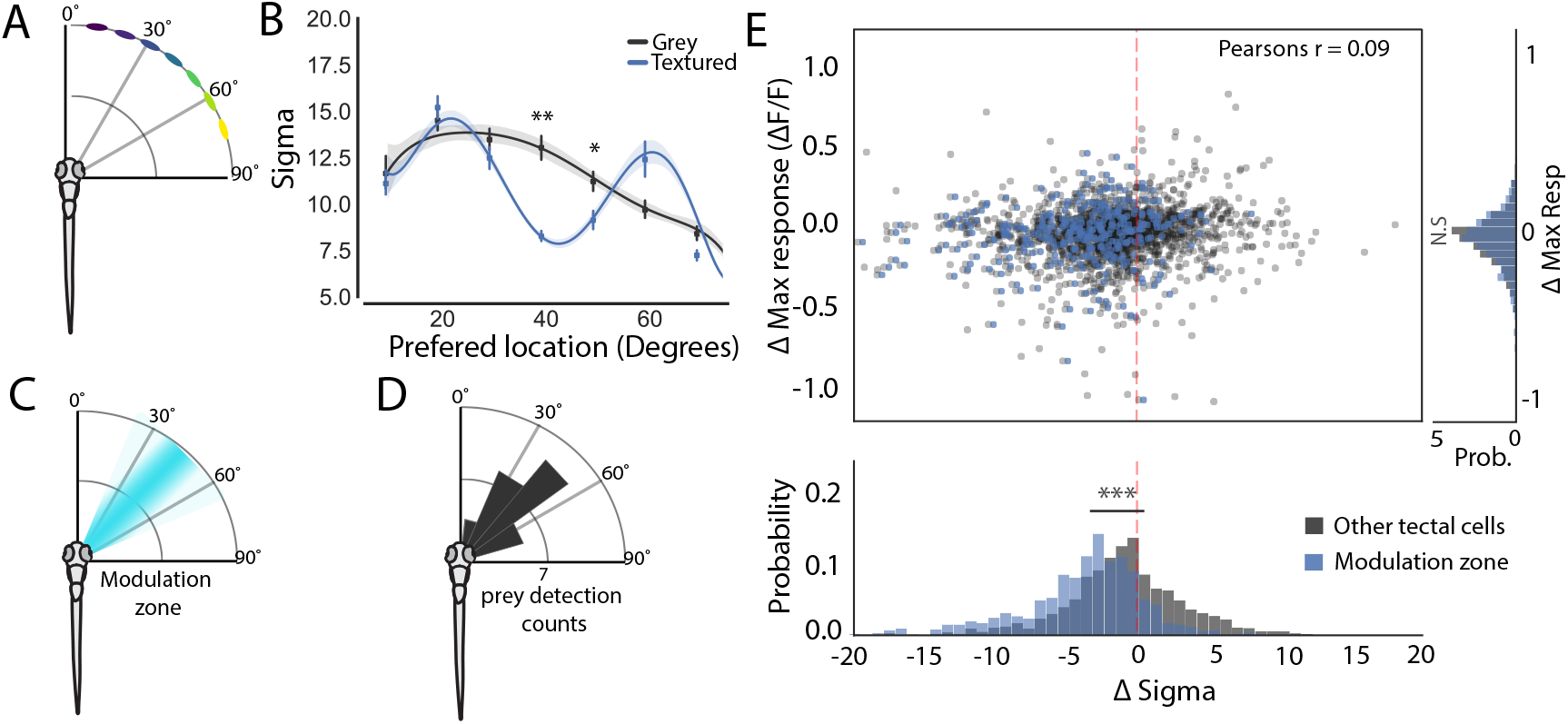
Contextual modulation takes place in a spatially restricted region of visual azimuth. (A) A schematic of stimulus location relative to the fish’s body axis (B) A plot of sigma against neuron’s preferred stimulus location for each fish, demonstrates that spatial tuning exhibits contextual sharpening only for stimuli presented between 35-50° of visual azimuth (40°: p =.002, 50°: p = 0.036, two-way ANOVA, T-tests with Bonferroni Correction) (C) Schematic showing the zone in visual space where contextual modulation occurs (D) This modulation zone corresponds to the area in visual space where hunting routines are preferentially triggered. (modified from ***Romano et al***. (***2015***)) (E) Top: A scatter plot showing that a neuron’s change in maximum response is not correlated with its change in it’s sigma (r=0.09, Pearsons). Cells within the modulation zone are highlighted in blue and were defined as cells which had a prefered tuning between 35-48°. Right: A histogram showing the difference in max response for neurons within the modulation zone and all other tuned neurons (p=.2). Bottom: A histogram showing the difference in delta sigma for neurons within the modulation zone and all other tuned neurons (p=10^-46^). * = p<.05, ** = p<.01, ** = p<.001.

One alternative explanation for the sharpening of tuning is that tectal responses to the prey-like stimuli are simply suppressed due to a reduction in contrast between the prey-like stimuli and the textured background. However, we found no correlation between each neuron’s change in maximum response (Δ Maximum response) and its change in (Δ sigma) (Fig. 2E). This suggests that the textured background sharpens spatial receptive fields without suppressing responses at preferred stimulus locations. In addition, visualising cells by their preferred location showed that cells within the contextually modulated area of visual space are reduced in their Δ sigma relative to all other cells (Fig. 2E). Together these results suggest that contextually modulated cells represent a distinct sub-type of cells within the tectum and which share a similar tuning preference.

Our results demonstrating that contextual modulation takes place within a defined region of visual azimuth, suggests that contextually modulated tectal neurons are localised to a topographically restricted region of the tectum since the tectum contains a retinotopic map of visual space (***Goodhill and Xu*** (***2005***)) (Fig. 3A). To map the tectal location of contextually modulated neurons, all imaged fish were transformed into a standard coordinate system (see Materials and Methods) (Fig. 3B). As expected, colour-coding these neurons according to their preferred stimulus location revealed an ordered topographic map of visual azimuth space along the anterior-posterior axis of the tectum (Fig. 3C). To understand how contextually modulated cells are distributed within the tectum, the location of highly modulated cells (cells with a Δ sigma < −5) were visualised as a density map over the tectum (Fig. 3D). This revealed that these cells tended to be grouped in the middle of the tectum’s anterior-posterior axis with a slight posterior bias. To quantify this topography the tectum was divided into zones along this axis and the mean sigma was calculated for each zone for each stimulus block. This showed that only neurons in the middle zones of the tectum showed reduced sigma’s in the textured block. This effect was also visible when mean Δ sigma was calculated for each segment (Fig. 3G), showing that the center of the tectum showed large negative changes in sigma that were not present at the tectal poles. Therefore, just as there is a modulation zone within visual space there is a corresponding region of the tectum where cells are preferentially modulated.

**Figure 3.**
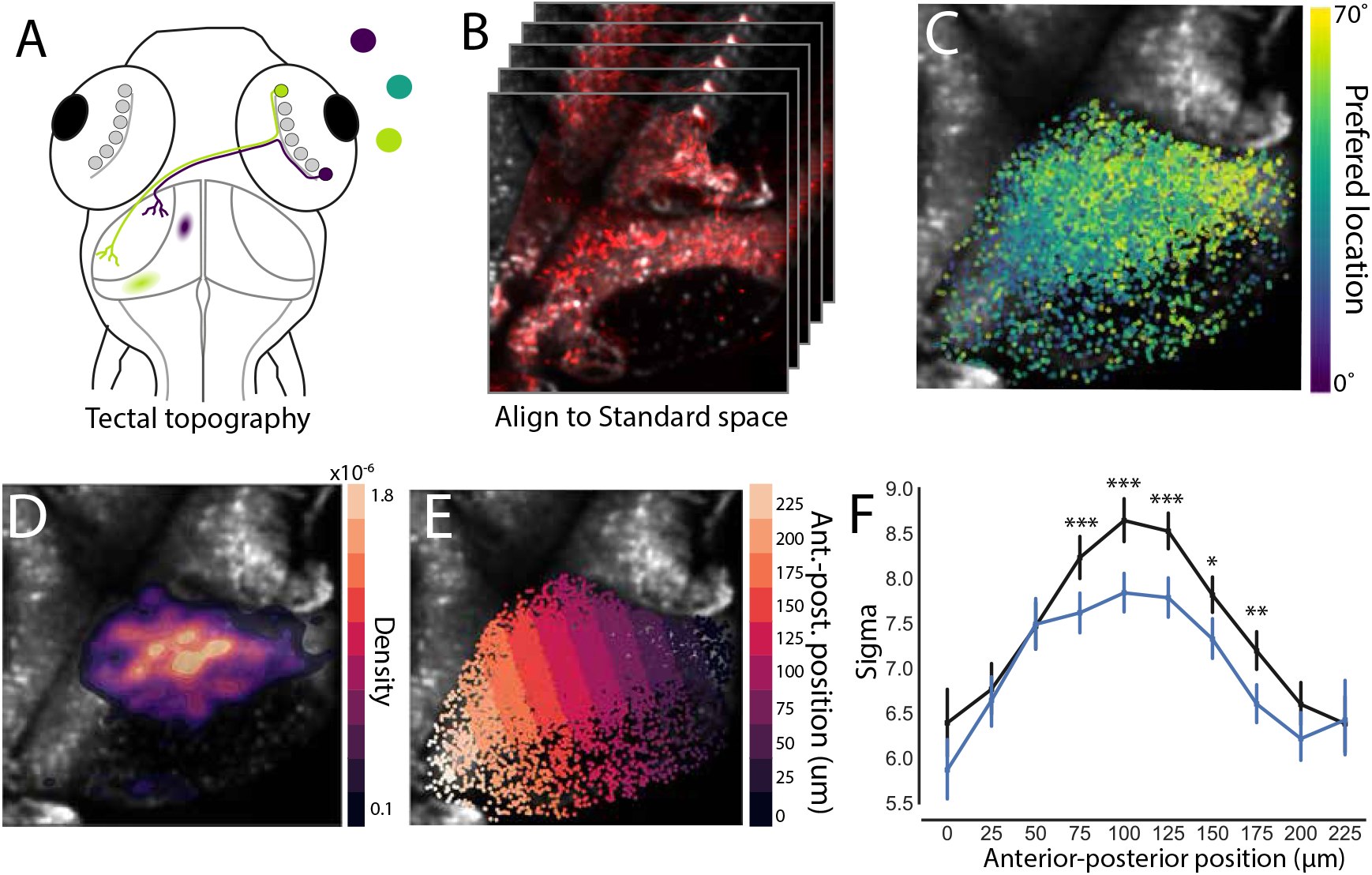
Modulated neurons are topographically distinct within the tectum. (A) A schematic detailing the topographic organisation of the tectum. Here retinal ganglion cells project out of the retina and make synapses in the neuropil of the contralateral hemisphere. They do this is a way that preserves a spatial map of visual space within the tectum with frontal visual space mapping onto the anterior portion of the tectum (purple), whereas rear visual space maps more posteriorly (lime green) (B) To assess the spatial arrangement of contextually modulated cells in the tectum a standard coordinate space was generated by aligning the functional imaging data to a high resolution stack of the tectum. (C) An overlay of cells in the tectum which have been colored by their tuning preference to demonstrate the topography of the tectum. (D) An overlay of a density heatmap showing the position of highly contextually modulated cells (Δ sigma < −5) within the tectum. (E) To quantify the position of contextually modulated cells the tectum was divided into bins along its anterior-posterior axis. (F) A plot of sigma values for each segment within the anterior-posterior axis for both textured and grey backgrounds (75*μ*m: p=10^-3^,100*μ*m: p=10^-6^,125*μ*m: p=10^-5^,150*μ*m: p=.02,175*μ*m: p=.002, two-way ANOVA, Bonferroni multiple comparison correction) (F) A plot showing Δ sigma by anterior-posterior segments. * = p<.05, ** = p<.01, ** = p<.001

In the visual cortex of mice, certain types of contextual modulation requires visual experience to develop (***Pecka et al***. (***2014***)). To test if this was also the case for zebrafish, larvae were reared from 0-7dpf either over a bed of gravel (GR), exposing them to natural visual features, or dark reared (DR) to deprive them of visual stimuli (Fig. 4A). To determine if a modulation zone was present in these fish the preferred location for each neurons was plotted against sigma value for both the textured and grey blocks. This revealed that regardless of rearing condition reduced sigma values were seen at 40° in the textured block, suggesting that a modulation zone was present in both rearing conditions. Furthermore, the average magnitude of Δ sigma was the same in both rearing conditions. These results show that development of circuits that support contextual modulation of spatial receptive fields in the zebrafish tectum is not dependent on visual experience.

**Figure 4.**
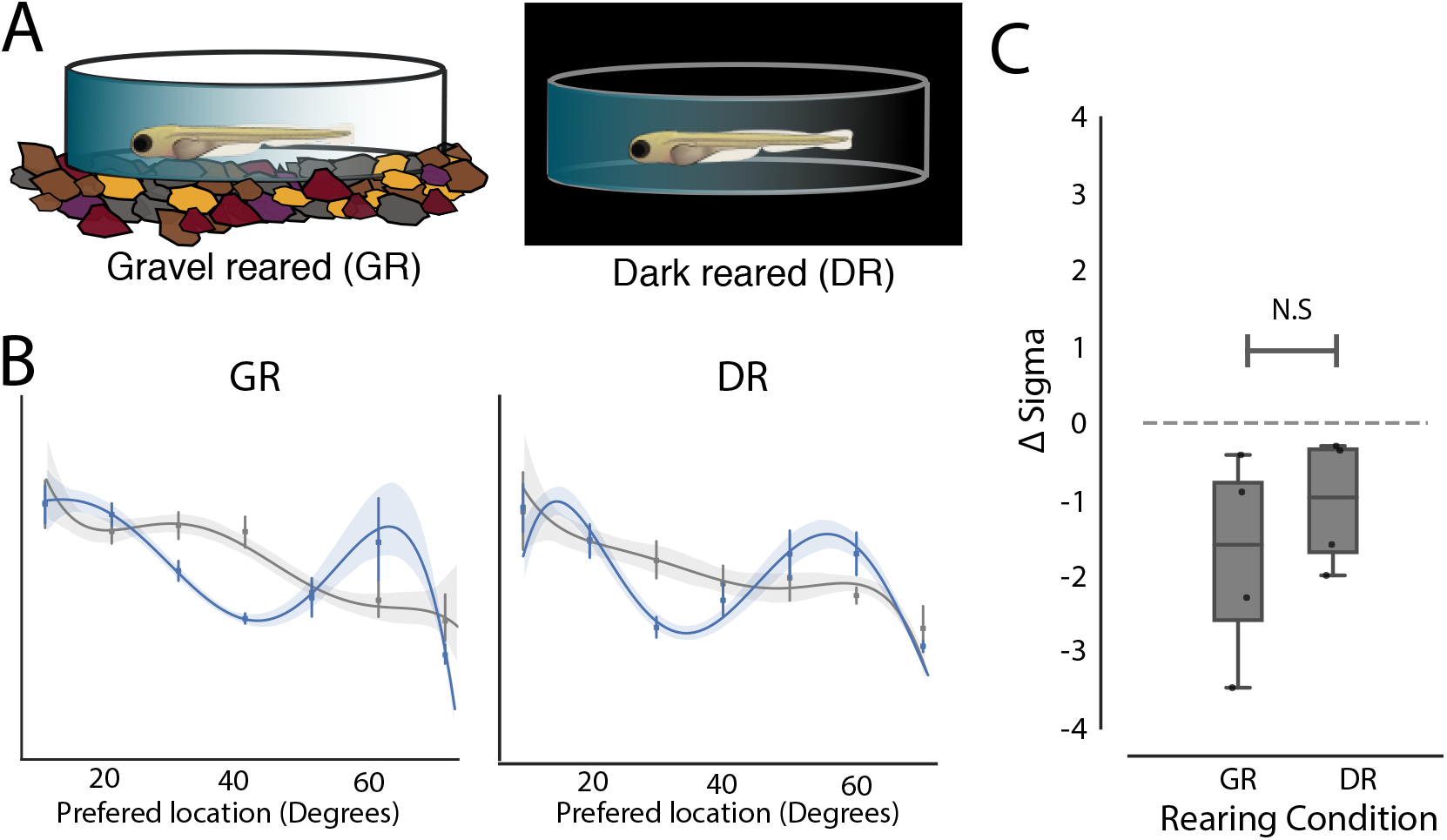
Visual experience has no effect on the development of contextual modulation in the optic tectum. (A) To alter the visual environment zebrafish were either raised over a bed of gravel (GR, n=4) or in complete darkness (DR, n=4). (B) Plot of cell’s preferred stimulus location against its sigma for both rearing conditions and in textured and grey stimulus blocks. A modulation zone can be seen in both conditions. (C) A boxplot showing no difference in delta sigma between background for GR and DR fish (p = 0.9, t-test).

## Discussion

Across multiple species it is well established that stimulating both a neurons RF and nRF with naturalistic stimuli can increase the selectivity of neurons for particular visual features (***Vinje and Gallant*** (***2000***); ***Haider et al***. (***2010***); ***Pecka et al***. (***2014***)). Likewise, in our study, we find a subset of tectal neurons become more sharply tuned to the position of prey-like stimuli when they are presented against a picture of gravel. This suggests that the objects in the nRF may be providing spatial information that reduces the uncertainty over the position of local objects. Therefore it is possible that neighbouring neurons with their receptive fields targeted to the contextually modulated neuron’s nRF may shape it’s spatial tuning through lateral GABAergic inhibition, as has been seen for other types of contextual modulation including those found in other teleost species (***Pecka et al***. (***2014***); ***Ben-Tov et al***. (***2015***)).

Interestingly, contextual modulation in the tectum was found not to be uniform, occurring in only a subset of tectal neurons. The receptive fields of of these neurons shared a tuning preference, constituting a modulation zone in visual space. This zone corresponds to a region in visual space where the presence of prey preferentially triggers the onset of hunting routines, characterised J-turns which orientate the larvae towards the prey (***Romano et al***. (***2015***); ***McElligott and O’Malley*** (***2005***); ***Bianco et al***. (***2011***)). Therefore it is possible that the sharpening of these receptive fields helps the fish to localise objects, such as prey, during the routines. Furthermore, we find the cell bodies of these neurons to be spatially clustered within the tectum, adding to a growing body of literature showing that tectal circuits show regional specialization which correlates with aspects of prey capture (***Zimmermann et al***. (***2018***); ***Förster et al***. (***2020***); ***Avitan et al***. (***2019***).

While contextual modulation in mouse V1 is strongly affected by natural scenes, this property is not present at eye opening. Instead, it requires experience of natural features to develop (***Pecka et al***. (***2014***)). In contrast, we found that contextual modulation in the optic tectum of zebrafish develops normally in fish that had been deprived of sensory experience. This is interesting because other visual features that have been found to require experience to form in other species have been found to be experience independent in fish (***Nikolaou et al***. (***2012***); ***Niell and Smith*** (***2005***); ***Gebhardt et al***. (***2019***)). This may reflect that fact that larvae begin to hunt at just 5 dpf. As a result, many aspects of develop may need to to be hardwired allowing for the rapid assembly of tectal circuits required for hunting (***Kutsarova et al***. (***2017***); ***Zhang et al***. (***2016***)). Therefore perhaps contextual modulation in the optic tectum has evolved ensure that fish are able to hunt at this young age.

Overall, our study represents the first description of the arrangement of contextually modulated cells in the visual system of a the larval zebrafish. We anticipate that this system will act as a useful model for understanding understanding the development of contextual modulation both in terms of circuit organisation and the genetic processes that are likely to drive its formation.

## Methods and Materials

### Animals and Rearing

Calcium imaging experiments were carried out in transgenic zebrafish with pan-neuronal expression of nuclear-localised GCaMP6s *Tg*(*HuC:H2B-GCaMP6s; casper*) (Ahrens lab, Janelia farm). All larvae were raised at 28.5°C in Danieau solution (58mM NaCl, 0.7 mM KCl, 0.4 mM MgSO4, 0.6 mM Ca(NO3)2, 5 mM HEPES, pH 7.6) and were exposed to a 14 hour ON/10 hour OFF light/dark cycle. Larvae were fed daily from 5 dpf using live rotifiers. To assess the impact of visual experience on the contextual modulation in the tectum, zebrafish were raised in either in total darkness (dark-reared, DR) or on a bed of gravel (gravel reared - GR) from 0-7 dpf. This work was approved by the local Animal Care and Use Committee (King’s College London), and was carried out in accordance with the Animals (Experimental Procedures) Act, 1986, under license from the United Kingdom Home Office.

### 2-photon Volumetric Calcium Imaging

Neural activity was monitored by imaging the calcium dynamics of between 500-1500 neurons in the tectal hemisphere that was contralateral to the eye recieving the visual stimulation with a custom built 2-photon microscope (Independent NeuroScience Services). Excitation was provided by a Mai Tai HP ultrafast Ti:Sapphire laser (Spectraphysics) tuned to 940nm. Laser power at the objective was kept below 18 mW for all fish. Emitted light was collected by a 16x, 1 NA water immersion objective (Nikon) and detected using a gallium arsenide phosphide detector (ThorLabs). Images (256 x 256 pixels) were acquired at a frame rate of 60Hz by scanning the laser in the x-axis with a resonant scanner and in the y-axis by a galvo-mirror. The focal plane was adjusted in 15*μ*m steps using a piezo lens holder (Physik Instrumente). This allowed for volumetric data consisting of 5 focal planes to be collected at a volume rate of 9.7Hz. Scanning and image acquisition were controlled by Scanimage Software (Vidrio Technologies). Each fish was imaged for 1 hour.

### The Visual Stimulation Setup

To record visually evoked responses within the tectum, 7 dpf zebrafish were mounted in 2% agarose within a custom built perspex cylindrical chamber. The fish was positioned so that its right eye faced a semi-circular screen covered in a grey diffusive filter and the chamber was filled with Danieau solution. This screen occupied 153° X 97° of visual azimuth and elevation respectively, and was positioned 20 mm away from the fish. Visual stimuli could then be projected onto this screen using a P2JR pico-projector (AAXA Tech). To avoid interference of the projected image with the signal collected by the detector, a red long-pass filter (Zeiss LP590 filter) was placed in front of the projector.

### Visual stimuli

Visual stimuli were generated using a custom C^++^ script written by Giovanni Dianna, Meyer lab. 5° black spots were presented at three different locations in visual azimuth separated by 10° intervals (10°, 20°, 30°, 40°, 50°, 60°, 70°). 0° was defined as midline directly in front of the fish. These spots moved with motion that resembled rotifer movement within a neighbourhood (5° radius) at a speed of 30°/S.

These dots were presented in two blocks which differed in the background against which they were projected. In one block the background was a picture of gravel (textured block) and the other it was simply a grey screen (grey block). In these blocks each spot was presented a total of 7 times per block in 5 second epochs, followed by 30 seconds of black screen. Importantly the movement of the dot was identical in each presentation. Both the order of the blocks and order of these spots within the blocks were pseudo-randomised. To prevent startling the fish, all dots faded in and out over the course of 1 second to minimise any startle effects that may be caused by sudden changes in the stimulus.

### Preprocessing of Calcium imaging data

Visually evoked functional imaging data was both aligned and segmented using the Suite2p Python package (https://mouseland.github.io/suite2p, ***Pachitariu et al***. (***2016***)). Only segments within the tectum with a probability > 0.5 of being a cell were used for further analysis.

To get smooth ΔF/F signals signals, free from imaging noise, the calcium signal was estimated from the raw fluroesence trace using the AR1 model contained within the OASIS package, with all parameters of the model being estimated from the data (***Friedrich et al***. (***2017***)). These were then used to calculate tuning profiles for each cell.

### Generating Tuning Profiles

To generate tuning profiles the max response to the stimulus was calculated for each repetition by first taking the max amplitude for each stimulus presentation. These amplitudes were then averaged for each stimulus location across repetitions, providing for each neuron a curve of the averaged max amplitude for each stimulus location. These coarse grained curves were then interpolated with a cubic spline at 5 degree intervals. These interpolated curves were then fitted with to a Gaussian function with a baseline offset using non-linear least squares. Initial parameters for fitting used the mean as the stimulus location eliciting response peak amplitude, and the initial value for standard deviation was varied from low to high values. The highest goodness of fit was selected (adjusted r2) was selected as the tuning profile for each neuron. This procedure was repeated twice for each cell, once for the grey block and once for the textured block. Only neurons with a goodness of fit greater that 0.9 in both blocks were used for further analysis. For these cells the mean of the Gaussian defined the preferred location whereas its standard deviation (sigma) quantifies the sharpness of tuning. By taking the difference in sigma (Δ sigma) for each neuron between the textured and grey blocks the change in the neurons tuning could be quantified. These values could then be visualised against each neurons preferred location to understand the distribution of contextually modulated cells in visual space.

### Calculating the topographic arrangement of contextually tectal neurons

To visualise topographic arrangement of contextually modulated cells in the tectum, cells from different fish needed to be transformed into a standard coordinate space. This required that the mean image for each functional imaging slice was aligned a reference 2-photon stack of the entire tectum hemisphere. In this reference stack the x and y coordinates were the same but 200 slices inz were taken ata resolution of 2*μ*m. Functional imaging data was aligned to this reference stack using the “SyN” method contained in the ANTsPy package. This method performs non-rigid alignment by applying both affine and deformable transformations and uses the mutual information between both stacks of images as an optimisation metric. The transformations from this alignment where the applied to the center point for each cell segment obtained using suite2p. This resulted in all neurons from each fish being put into a standardised coordinate space which could then be used to look localise cells within the tectum whose tuning was modulated by context.

Once in this space principal component analysis was applied to the x-y positions of the tectal neuron segments. The first principle component spanned the major axis of the tectum, corresponding to the anterior-posterior axis. This axis could then be divided into bins and the sigma values for each bin could be obtained.

### Sample sizes and statistical analysis

Multiple comparisons were first tested with and either one or two-way ANOVA depending on the number of factors being compared. If significance was reached post-hoc t-tests with multiple comparisons were used with the method of correction. All tests and significance are reported in the figure legends throughout. All sample sizes are similar to those typically used in the zebrafish imaging field.

## Acknowledgments

We would like to thank Misha Ahrens for the zebrafish lines. We would also like to acknowledge Juan Burrone, Matt Grubb and Adil Khan for their well constructed criticism when writing the manuscript

